# Multi-omic signatures identify pan-cancer classes of tumors beyond tissue of origin

**DOI:** 10.1101/806323

**Authors:** Agustin Gonzalez-Reymundez, Ana I. Vazquez

**Affiliations:** Department of Epidemiology and Biostatistics, Michigan State University, MI, USA; Institute for Quantitative Health Science and Engineering (IQ), Michigan State University, MI, USA

## Abstract

Despite recent advances in treatment, cancer continues to be one of the most lethal human maladies. One of the challenges of cancer treatment is the extreme diversity among seemingly identical tumors: while some tumors may have good prognosis and are treatable, others are quite aggressive, and may lack of effective therapies. Most of this variability comes from wide-spread mutations and epigenetic alterations. Using a novel omic-integration method, we have exploited this molecular information to re-classify tumors beyond the constraints of cell type. Eight novel tumor groups (C1-8) emerged, characterized by unique cancer signatures. C3 had better prognosis, genome stability, and immune infiltration. C2 and C5 had higher genome instability and poorer clinical outcomes. Remaining clusters were characterized by worse outcomes, along with higher genome instability. C1, C7, and C8 were upregulated for cellular and mitochondrial translation, and relatively low proliferation. C6 and C4 were also downregulated for cellular and mitochondrial translation, and had high proliferation rates. C4 was represented by copy losses on chromosome 6, and had the highest number of metastatic samples. C8 was characterized by copy losses on chromosome 11, having also the lowest lymphocytic infiltration rate. C6 had the lowest natural killer infiltration rate and was represented by copy gains of genes in chromosome 11. C7 was represented by copy gains on chromosome 6, and had the highest upregulation in mitochondrial translation. We believe that, since molecularly alike tumors could respond similarly to treatment, our results could inform therapeutic action.

**Significance:** Cancer has been traditionally studied as a family of different diseases from different anatomical sites. Nevertheless, regardless of the tissue of origin, cancer can be characterized by molecular alterations on mechanisms controlling cell fate and progression. In this study, we integrate 33 cancer types and show the existence of eight clusters with unique genomic signatures and clinical characteristics, beyond the site of origin of the tumor. The study and treatment of cancer, based on predominant molecular features, rather than site of origin, can potentially aid in the discovery of novel therapeutic alternatives.

## Introduction

In spite of recent advances that have improved the treatment of cancer, it continues to reign as one of the most lethal human diseases. More than 1,700,000 new cancer cases and more than 60,000 deaths are estimated to occur in the year 2019, in the United States alone^1^. Cancer can be considered a highly heterogeneous set of diseases: while some tumors may have a good prognosis and are treatable, others are quite aggressive, lethal, or may not have a standard of care^2–4^. Cancer can also defy standard classification: a well classified tumor may not respond to standard therapy, as expected, and may behave as a different cancer type^5–7^. Fortunately, with the advances of sequencing technologies, data has become available for research as never before. The Cancer Genome Atlas (TCGA), for instance, offers clinical and omic (e.g. genomic, transcriptiomic, and epigenenomic data) information from more than 10,000 tumors across 33 different cancer types^8^. Much of this omic data has the potential to enable us to classify tumors and to explain the striking variation observed in clinical phenotypes ^9–12^.

Omic integration has been successfully applied in previous classification efforts^13–16^. These classifications have highlighted how molecular groups of tumors highly agree with human cell types. Alternatively, we hypothesize the existence of internal subtypes hidden by cell type and tissue characteristics influencing cell behavior. These subtypes could be distinguished by molecular alterations unlocking cancerous cell-transformation events. To test this hypothesis, we have developed a statistical framework that summarizes omic patterns in main axes of variation describing the molecular variability among tumors. Key features characterizing each axis (i.e. features contributing the most to inter-tumor variability) are retained, while irrelevant ones are filtered. Retained features are then used to cluster tumors by molecular similarities and find specific molecular features representing each group.

Here we show that, after removing all tissue-specific effects, the cancer signal immediately emerges. The new molecular aggrupation, emphasizing on shared tumor biology, has the potential of providing new insights of cancer phenotypes. We expect this novel classification to contribute to the treatment of tumors without a current standard of care, by for example, borrowing therapies from molecularly similar cases.

## Results

Signal coming from tissue and cell type strongly influence a naïve initial classification of tumors across cancer types. We performed omic integration based on penalized matrix factorization, in order to remove tissue effects, and seek out a re-classification of tumors based on subtler omic patterns. Our method can be illustrated in four steps (Figure 1, Materials and Methods). *Step 1* consists of applying sparse Singular Value Decomposition (sSVD) to an extended omic matrix ***X***, obtained from concatenating a series of scaled and normalized omic blocks for the same subjects. Briefly, the major axes of variation across tumors (i.e. left principal components, or scores) and the matching features ‘activities’ (i.e. the right principal components, or loadings) of ***X*** are found. Sparsity is then imposed on the activity values, so features with minor influence over the variability among tumors, are removed. *Step 2* consists of identifying what features (expression of genes, methylation intensities, copy gains/losses) influence these axes the most (i.e. features not removed by sSVD) and mapping them onto genes and functional classes (e.g. pathways, ontologies, targets of micro RNA). *Step 3* involves the identification of local clusters of tumors, following Taskensen et al. (2016). *Step 4* involves the characterization of clusters in terms of molecular (e.g. genes, pathways, complexes, etc.) and clinical (e.g. survival probability, immune infiltration, etc.) information, distinguishing each cluster from the rest.

**Figure 1:**
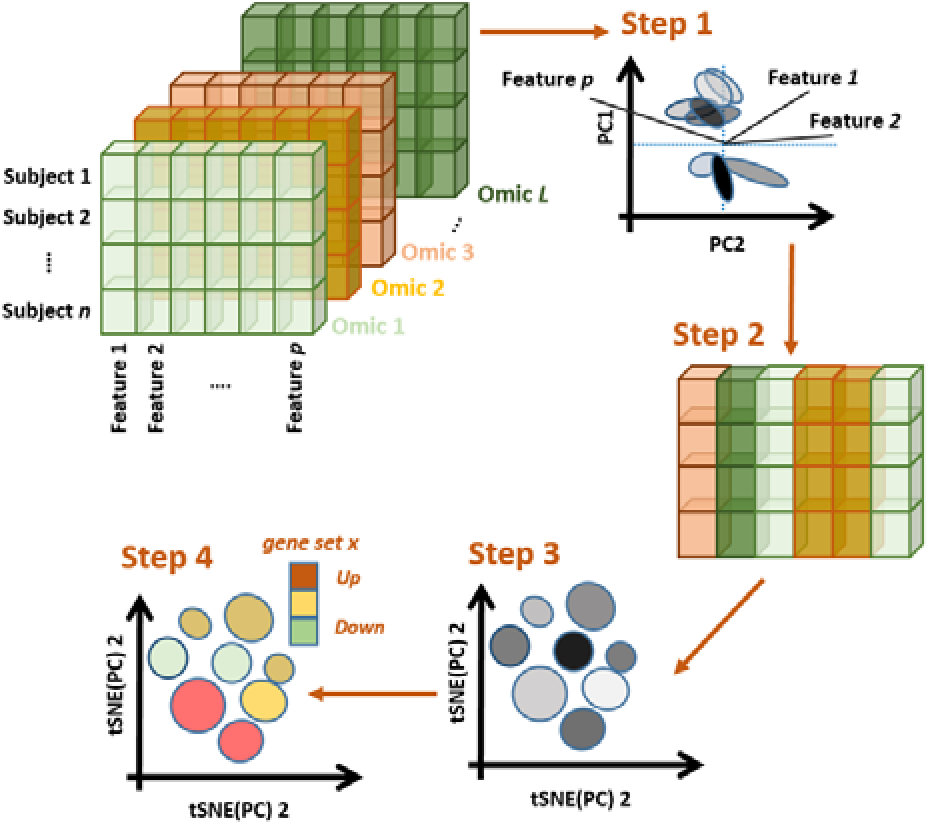
Omic integration and features selection method. *Step 1)* Singular value decomposition of a concatenated list of omic blocks and identification of major axes of variation. *Step 2)* Identification of omic features (expression of genes, methylation intensities, copy gains/losses) influencing the axes and mapping them onto genes and functional classes (e.g. pathways, ontologies, targets of micro RNA). *Step 3)* Mapping major axes of variation via tSNE and cluster definition by DBSCAN. *Step 4)* Phenotypic characterization of each cluster of subjects.

Using samples from 33 different cancer types provided by The Cancer Genome Atlas (TCGA), and accompanying information from whole genome profiles of gene expression (GE), DNA methylation (METH) and copy number variant alterations (CNV), we re-classified tumors based on molecular similarities between the three omics. This was done by first removing the non-cancer systematic effects of tissue via multiplication of ***X*** by a linear transformation (see Materials and Methods section).

### Data description

The data, including information of sample size and type of sample (i.e. from normal, metastatic, or primary tissue), demographics (age, sex, and ethnicity) and survival information (overall survival status and times), are summarized in Table 1. Omic data included information for gene expression (**GE**, as standardized log of RNAseq data for 20,319 genes), methylation (**METH**, as standardized M-values summarized at the level of 28,241 CpG islands), and copy number variants (**CNV**, as standardized log of copy/gain intensity summarized at the level of 11,552 genes).

**Table 1:** Data description by cancer type after quality control. Tumor samples are described by cancer type (TCGA **Codes** and cancer name), in terms of relative sample size (**n**), percent of females (**F%**), ethnicities (percent of non-Hispanic Whites, Afro-descendants, and Asians), **Age** (at the moment of diagnosis, in years), type of sample (**TS%**, as percent of normal –**N**- and metastatic –**M**- samples), and survival (**Surv**, as expected time to 50% survival, in years). **Age** and **Surv** are represented by median values, with first and third quartiles as measurements of dispersion. Data corresponded to the alignment and intersection of all samples with information of gene expression (**GE**), methylation (**METH**), and copy number variants (**CNV**).

| Code | Cancer type | n | F% | Ethnicity %* |  |  | Age | TS% |  | Surv |
| --- | --- | --- | --- | --- | --- | --- | --- | --- | --- | --- |
|  |  |  |  | AD | W | A |  | N | M |  |
| **ACC** | Adrenocortical carcinoma | 23 | 61 | 0 | 100 | 0 | 48 (35-57) | 0 | 0 | 6.6 (2.5-6.6) |
| **BLCA** | Bladder urothelial carcinoma | 271 | 99 | 13 | 80 | 7 | 58 (49-66) | 1 | 0 | 3.0 (1.2-3.0) |
| **BRCA** | Breast invasive carcinoma | 639 | 69 | 18 | 75 | 7 | 58 (46-71) | 7 | 0 | 10.2 (6.5-10.2) |
| **CESC** | Cervical squamous cell carcinoma and endocervical adenocarcinoma | 234 | 25 | 8 | 78 | 14 | 60 (53-69) | 1 | 1 | 11.2 (3.1-11.2) |
| **CHOL** | Cholangiocarcinoma | 12 | 36 | 0 | 100 | 0 | 55 (46-67) | 75 | 0 | 1.7 (0.7-5.3) |
| **COAD** | Colon adenocarcinoma | 264 | 36 | 12 | 79 | 9 | 58 (41-66) | 7 | 0 | 8.3 (3.6-8.3) |
| **DLBC** | Lymphoid Neoplasm Diffuse Large B-cell | 26 | 54 | 19 | 81 | 0 | 60 (54-63) | 0 | 0 | 17.6 (17.6-17.6) |
| **ESCA** | Esophageal carcinoma | 134 | 60 | 12 | 88 | 0 | 68 (59-73) | 2 | 0 | 2.3 (1.1-4.4) |
| **GBM** | Glioblastoma multiforme | 49 | 23 | 12 | 78 | 10 | 66 (60-73) | 0 | 0 | 0.9 (0.4-1.2) |
| **HNSC** | Head and Neck squamous cell | 89 | 48 | 8 | 91 | 1 | 61 (59-71) | 1 | 0 | 5.9 (1.2-5.9) |
| **KICH** | Kidney chromophobe | 2 | 0 | 0 | 100 | 0 | 52 (50-54) | 0 | 0 | --** |
| **KIRC** | Kidney renal clear cell carcinoma | 43 | 51 | 2 | 91 | 7 | 67 (62-75) | 0 | 0 | 7.5 (7.5-7.5) |
| **KIRP** | Kidney renal papillary cell carcinoma | 37 | 62 | 20 | 80 | 0 | 65 (59-72) | 0 | 0 | -- |
| **LAML** | Acute myeloid leukemia | 28 | 0 | 0 | 94 | 6 | 60 (57-67) | 0 | 0 | -- |
| **LGG** | Brain lower grade glioma | 93 | 42 | 11 | 88 | 1 | 70 (62-75) | 0 | 0 | 9.5(3.1-12.2) |
| **LIHC** | Liver hepatocellular carcinoma | 62 | 25 | 8 | 92 | 0 | 69 (61-74) | 13 | 0 | 4.6 (1.6-8.6) |
| **LUAD** | Lung adenocarcinoma | 381 | 29 | 6 | 90 | 5 | 66 (59-72) | 4 | 0 | 4.2 (2.1-9.2) |
| **LUSC** | Lung squamous cell carcinoma | 289 | 28 | 9 | 89 | 2 | 57 (46-64) | 0 | 0 | 4.7 (1.8-10.5) |
| **MESO** | Mesothelioma | 68 | 0 | 7 | 93 | 0 | 60 (53-66) | 0 | 0 | 1.6 (0.9-2.4) |
| **OV** | Ovarian serous cystadenocarcinoma | 5 | 0 | 0 | 100 | 0 | 60 (55-61) | 0 | 0 | 2.9 (2.9-2.9) |
| **PAAD** | Pancreatic adenocarcinoma | 151 | 24 | 4 | 76 | 20 | 67 (60-74) | 3 | 0 | 1.6 (1.0-4.1) |
| **PCPG** | Pheochromocytoma and paraganglioma | 144 | 0 | 0 | 100 | 0 | 61 (56-65) | 0 | 1 | -- |
| **PRAD** | Prostate adenocarcinoma | 490 | 36 | 5 | 94 | 1 | 62 (54-70) | 6 | 0 | 9.6 (9.6-9.6) |
| **READ** | Rectum adenocarcinoma | 83 | 42 | 0 | 85 | 15 | 63 (54-73) | 2 | 0 | 3.9 (3.9-3.9) |
| **SARC** | Sarcoma | 181 | 41 | 0 | 100 | 0 | 58 (46-69) | 0 | 1 | 6.7 (3.1-6.7) |
| **SKCM** | Skin cutaneous melanoma | 378 | 85 | 15 | 83 | 2 | 61 (50-70) | 0 | 75 | 7.4 (2.6-20.1) |
| **STAD** | Stomach adenocarcinoma | 263 | 37 | 4 | 70 | 25 | 67 (58-73) | 0 | 0 | 4.6 (1.3-4.6) |
| **TGCT** | Testicular germ cell tumors | 134 | 0 | 4 | 92 | 4 | 31 (26-37) | 0 | 0 | -- |
| **THCA** | Thyroid carcinoma | 501 | 73 | 6 | 80 | 13 | 46 (35-58) | 8 | 1 | -- |
| **THYM** | Thymoma | 106 | 45 | 6 | 85 | 9 | 58 (48-68) | 1 | 0 | 9.6 (9.6-9.6) |
| **UCEC** | Uterine corpus endometrial carcinoma | 146 | 100 | 43 | 57 | 0 | 65 (57-72) | 14 | 0 | 9.2 (3.6-9.2) |
| **UCS** | Uterine carcinosarcoma | 4 | 100 | 0 | 75 | 25 | 63 (54-74) | 0 | 0 | 1.4 (0.3-2.2) |
| **UVM** | Uveal melanoma | 78 | 45 | 0 | 100 | 0 | 62 (51-74) | 0 | 0 | 3.8 (2.4-3.8) |

\*: Only the three most abundant ethnicities in the data set were considered to calculate the percent.
\*\*: Survival quantiles for cancer types with less than five death events were not calculated.

The first 50 main axes of variations of the extended omics matrix (selected by clear bend in the scree plot of Eigen-values – see Material and Methods). The projection of the 50 axes onto two dimensions is shown in Fig. S1. As expected, cell-of-origin effects dominate the clustering of tumors at a pan-cancer level, with clusters enriched by previously reported pan-cancer clusters (e.g. collection of gastric cancer, gliomas, kidney and squamous tumors), types, and subtypes (e.g. Luminal and Basal breast tumors), and single cancer types (e.g. Thyroid carcinoma, Prostate adenocarcinoma, etc.).

### Re-classification of pan-cancer tumors based on similarities between omics after removing tissue specific signals

Once tissue signal was identified, it was removed from the extended omic matrix. Next, sparsity constraints were imposed on the omic features in order to zero-out the features with irrelevant contribution to axes of variation and cluster formation. The selected features (i.e. with non-zero effects) across the three omics corresponded with the 18^th^, 25^th^, 33^th^, and 38^th^ axes (sorted from more to less variance explained) and mapped onto a total of 1200 genes. The cluster identification and projection onto two dimensions revealed eight classes (Figure 2). As a consequence of removing the effects of tissue localization, all clusters were formed by samples coming from multiple cancer types. Some clusters differed statistically from their cancer types composition (Table 2). However, all cancer types overlapped with more than one cluster (Fig. 2; Table 2, bottom). Furthermore, this overlap was not influenced by previously reported subtypes (Fig. S2).

**Table 2:**
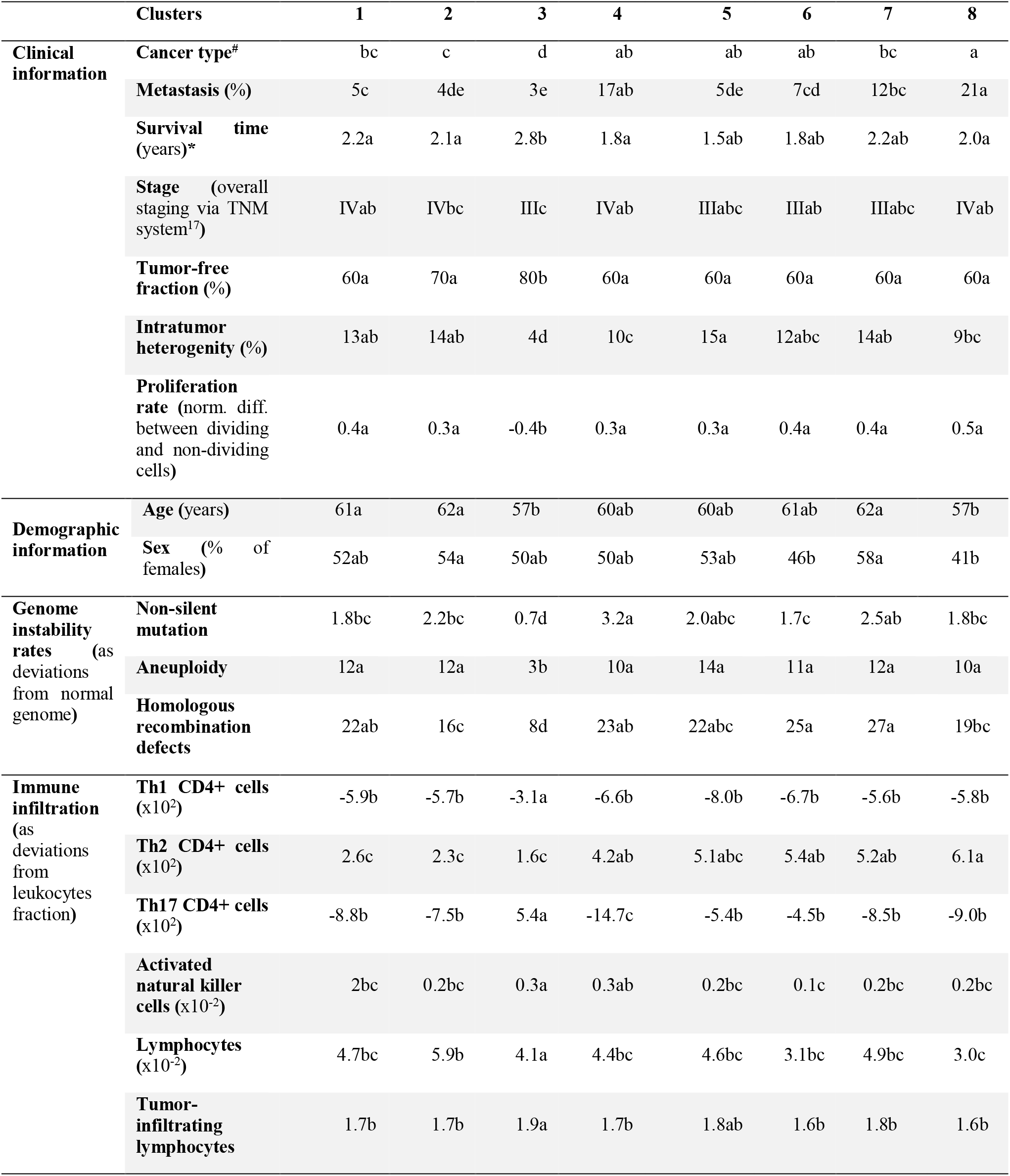
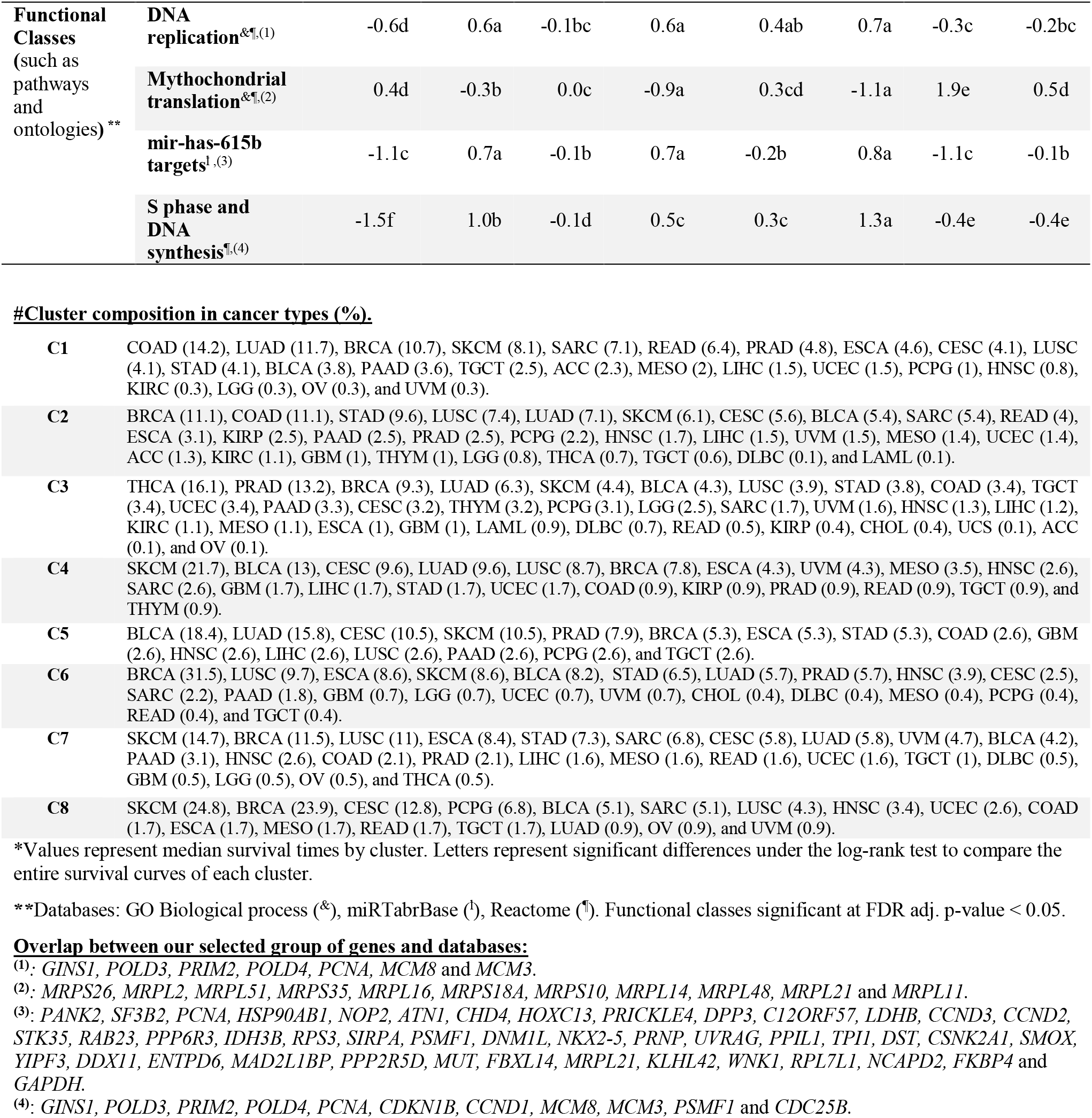
Characterization of pan-cancer clusters of tumors after removing tissue effects. The clusters produced by integration of whole-genome profiles of gene expression (GE), copy number variants (CNV), and DNA methylation (METH) were characterized in terms of clinical, demographic, immune and molecular information. The table shows those variables with significant differences in at least one cluster. For each variable, different letters represent significant differences between clusters.

**Figure 2:**
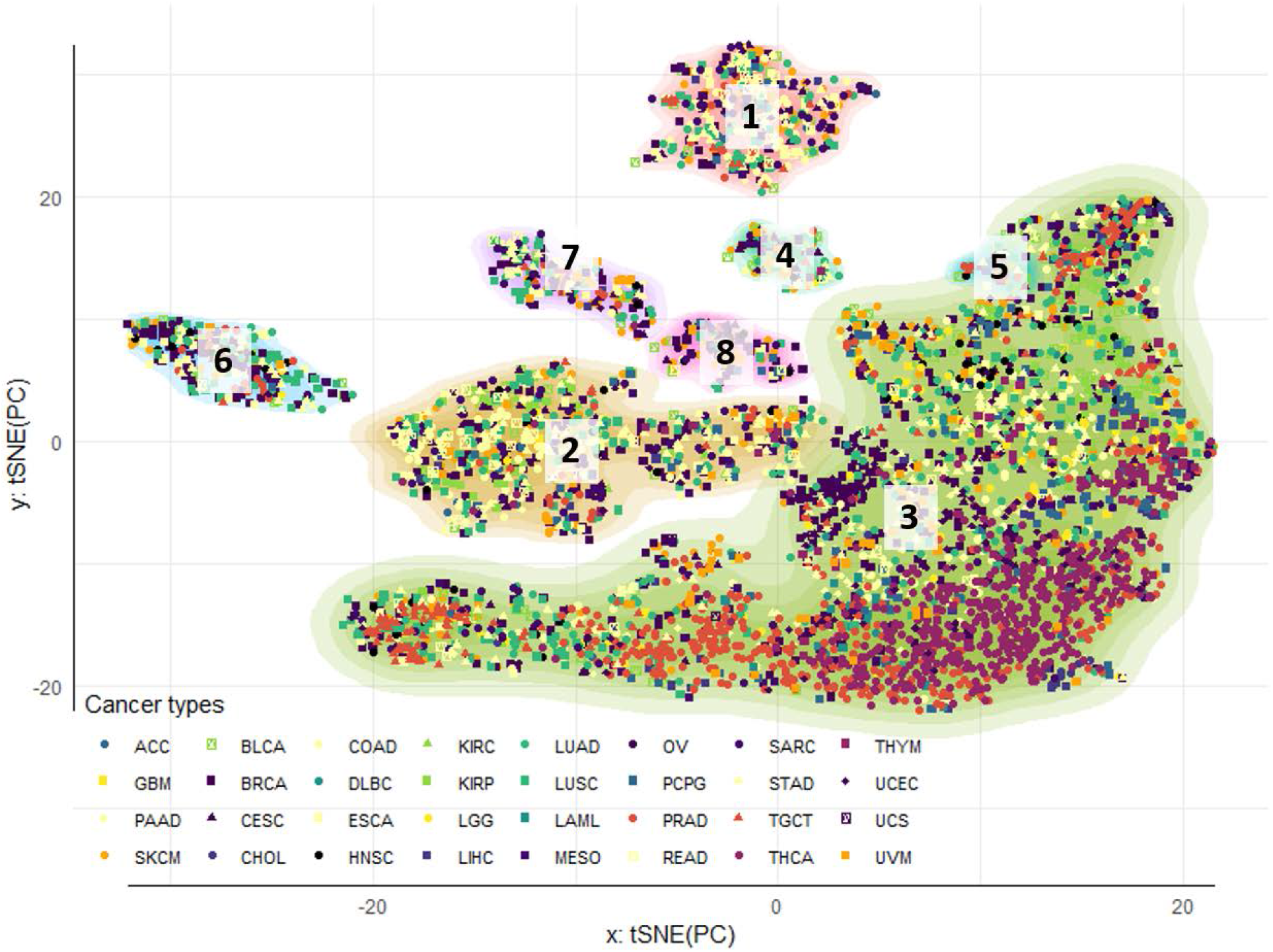
Pan-cancer clustering of tumor samples: tissue effects correction a selection of omic features. Tumor clusters were obtained by sequential application of tSNE and DBSCAN algorithm for 5,408 samples across 33 cancer types. The contours reflect cluster membership, and the points’ colors and shapes represent similar anatomical site and cancer type, respectively. The two-dimensional tSNE projection was obtained from the four deep principal axes of the extended omic matrix projected outside the tissue specific effects, after performing sSVD and removing the first two axes. After re-classifying tumors, the few samples coming from Kidney chromophobe tumors (KICH) did not map in any of the eight clusters obtained.

### Clinical and demographical characterization of tumor clusters

Clusters differed statistically in terms of patient age (with Cluster 3 and 8 containing samples from slightly younger patients) and sex (with Clusters 2 and 7 having significantly more females than Cluster 8, due to their slightly higher composition of gynecological cancers) (Table 2). None of the clusters were significantly associated with ethnicity (Table 2).

The most notorious distinctions between clusters were their differences in prognosis and severity traits (Fig. S3). Cluster 3 (the largest cluster in Fig. 2) was distinguished by better prognosis/less severity cancer than the remaining clusters, followed by Clusters 2, 5, 6 and 7. Clusters 4 and 8 were in general the ones with worst prognosis and more aggressive tumors (Table 2). Cluster 3 was also the one with fewest metastatic samples (Fig. S4), higher survival rates, highest tumor-free fraction, lowest stage, lowest intra-tumor heterogeneity (ITH, that estimates the fraction of subclonal and clonal genomes in each sample^18^), and lowest proliferation (Table 2, Fig S3). By comparison, Clusters 4 and 8 had significantly more metastatic samples than Cluster 3. Cluster 8 had also higher ITH rates than Cluster 3. The highest ITH rates were found in Cluster 5.

Cluster 3 had also the lowest rates of non-silent mutations, aneuploidies, and homologous recombination dysfunction (HRD). The remaining clusters were very similar in terms of genome instability indicators, except for Cluster 2. This cluster had significantly higher rates of HRD than Cluster 3, but significantly lower rates than every other cluster (Table 2). In terms of immune infiltration, Cluster 3 was characterized by the highest rates of tumor suppressive immune cells and tumor infiltrating lymphocytes (Table 2). In addition, Cluster 6 had the lowest infiltration of activated natural killer (ANK) cells. Cluster 8 had also the lowest lymphocytic and highest Th2 CD4+ infiltrations, respectively (Table 2).

### Gene signatures characterizing tumor clusters

The clusters were also characterized by distinct sets of omic features, significantly enriched for functions involved in cell cycle (DNA replication, DNA synthesis, and targets of hsa-mir-615-b, a micro RNA involved in cell proliferation) and mitochondrial translation (initiation, elongation, and termination) (Table 2). To study the pairwise differences across clusters, these gene sets were projected onto scores for each gene, as linear combinations between the features’ values mapping onto the gene (i.e. its expression, methylation, and copy number values) and their corresponding activities (i.e. the features effects arising from the sparsity constraints) (see Materials and Methods section). In general, Cluster 3 was characterized by intermediate values of these scores, while the remaining clusters were characterized by higher (i.e. gene set with higher expression than Cluster 3) or lower (gene sets with lower expression than in Cluster 3) gene set scores. Clusters 2, 4, and 6 had significantly higher scores for cell proliferation, and significantly lower for mitochondrial translation. Clusters 1, 7 and 8, on the other hand, had significantly lower scores of proliferation and higher for mitochondrial translation.

Sparse factorization of the extended omic matrix resulted in the selection of features mapping onto 1200 genes. From this list, 441 genes were significantly different in at least one cluster. These results were obtained by a series of analyses of variance (ANOVAs), using the scores of each gene as response variables and clusters as explanatory variables. This list included 34 validated cancer genes, including oncogenes (*ERC1*, *HSP90AB1*, *NUMA1*, *PPFIBP1*, *ZNF384*, *CHD4*, *KRAS*, *HIST1H3B*, *CCND1*, *CCND2*, *PIM1*, *CCND3*, *HMGA1*, *HOXC11*, *HOXC13*, *KDM5A*, *SRSF3*, *TFEB*), tumor suppressors (*FANCE*, *CDKN1B*, *ASXL1*, *ETNK1*) and fusion-proteins (*ERC1*, *HSP90AB1*, *NUMA1*, *PPFIBP1*, *ZNF384*). Many of the genes additionally mapped onto known transcription factors (including: *KDM5A*, *RELA*, *SRF*, *CTBP2*, *FOXA2, NONOG, FOLSL1, TEAD4*, and *FOXM1*) and some of their targets (Fig. S5). However, the expressions of TFs and their targets were not significantly correlated within or between clusters (Fig. S5), suggesting mechanisms of control of the gene expression other than TFs regulation.

We then interrogated all pair-wise comparisons between the scores of each one of the 441 significant genes using Tukey tests (Supplementary Table S2). We identified a subgroup of 123 significant genes that distinguished each cluster from the rest (for example, *POLH* had significantly higher scores in Cluster 4 than in every other cluster). The genes characterizing each individual cluster were then used to define signatures. With this criterion, only Clusters 1, 4, 6, 7, and 8 were characterized by distinct signatures of 57, 4, 23, 24, and 15 genes each, respectively. Since the gene scores are combinations of omic features, we looked at the gene expression in each signature and the potential role of copy numbers and methylation in regulating it (Figures 3–4).

**Figure 3:**
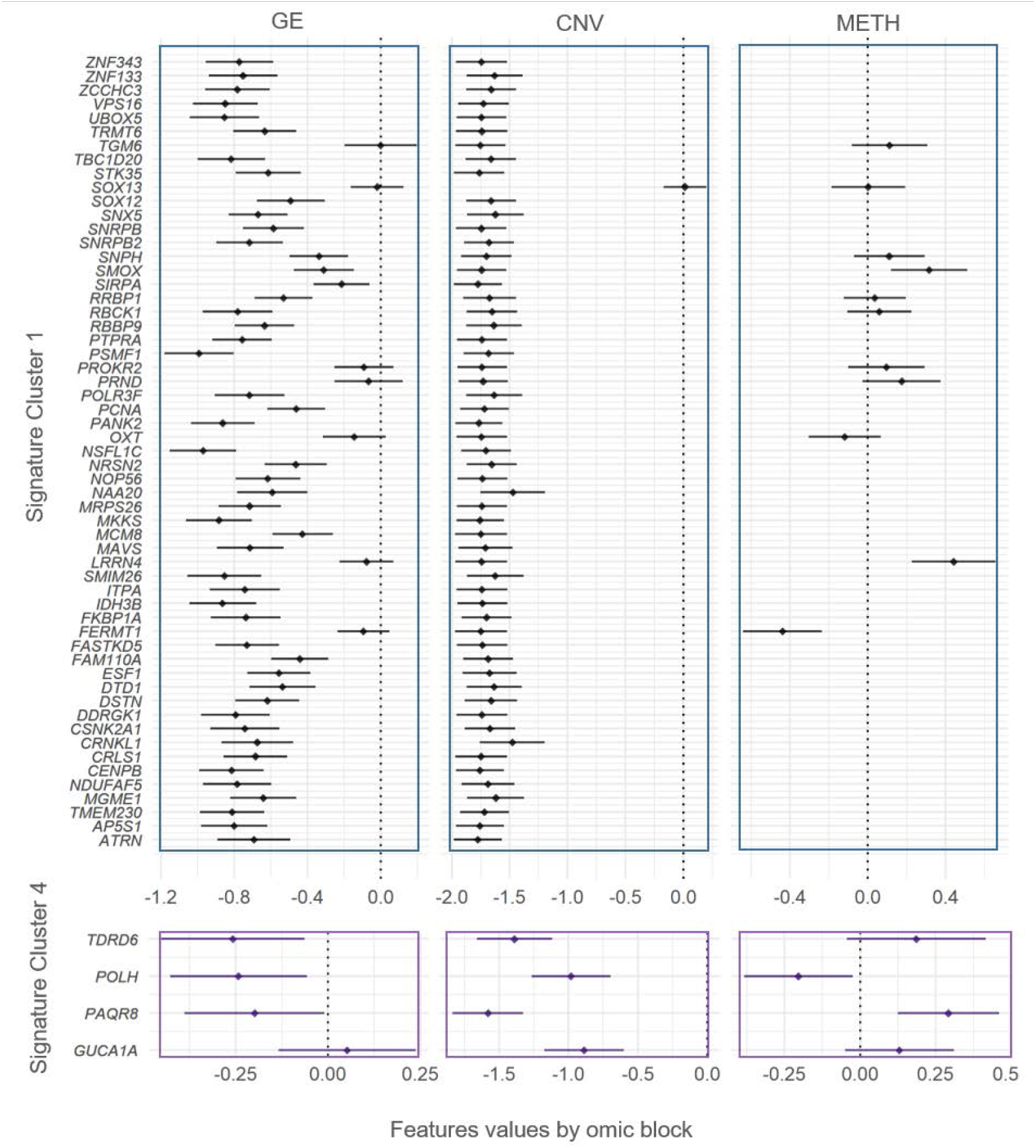
Gene signatures for Clusters 1 and 4 in terms of gene expression, copy number variation, and methylation. The genes significantly de-regulated exclusive of Clusters 1 and 4 were used to define signatures (y-axis). The features values (x-axis) of each gene are separated in gene expression (GE, first column of panels), copy number variants (CNV, second column of panels), and DNA methylation (METH, third column of panels), and summarized by Bonferroni confidence intervals (adjusting for all the 441 significant genes in at least one cluster). Dots represent the average of features values across samples.

**Figure 4:**
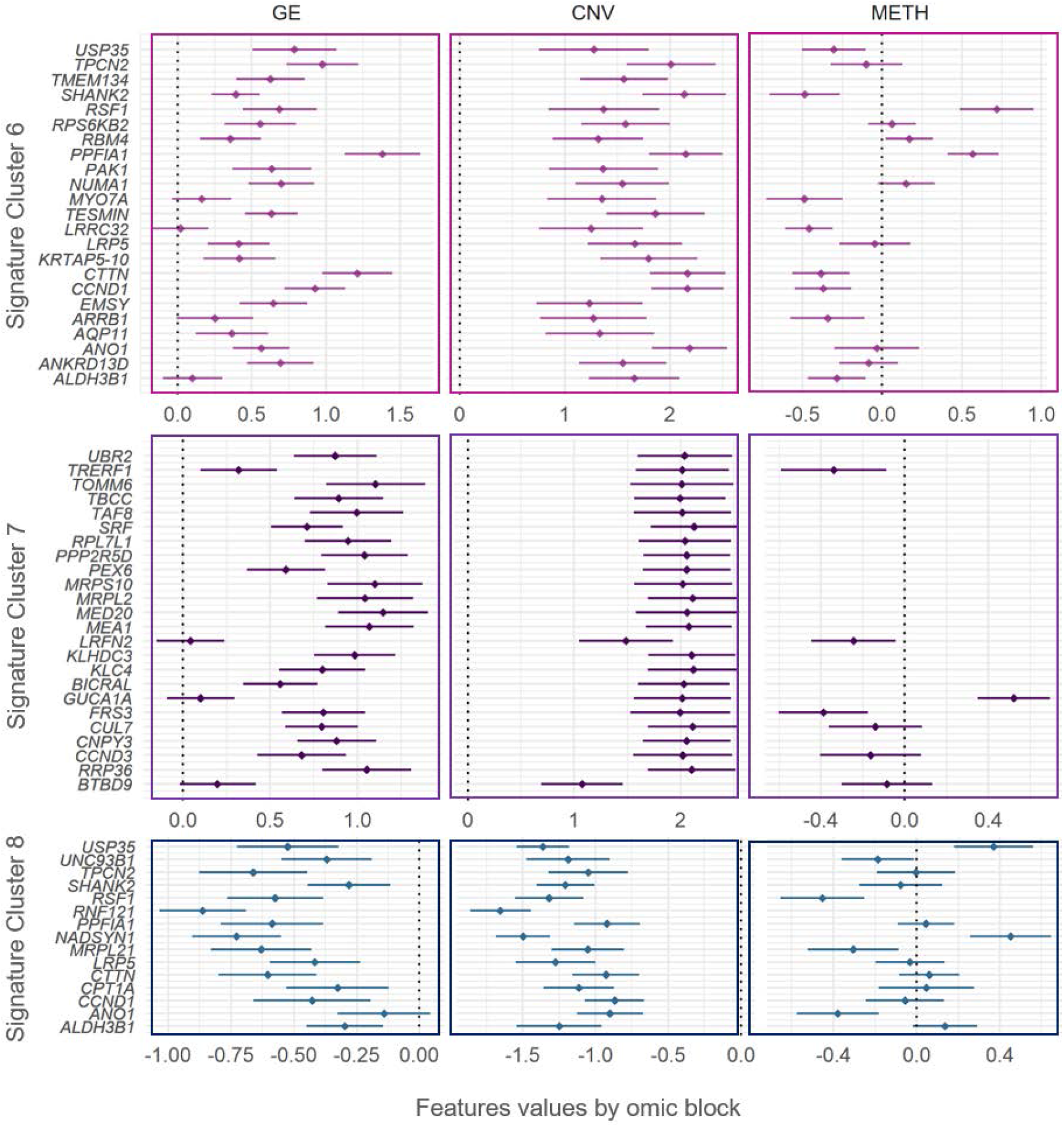
Gene signatures for Clusters 6, 7 and 8 in terms of gene expression, copy number variation, and methylation. The genes significantly de-regulated exclusively in Clusters 6, 7 and 8 were used to define signatures (y-axis). The features values (x-axis) of each gene are separated in gene expression (GE, first column of panels), copy number variants (CNV, second column of panels), and DNA methylation (METH, third column of panels), and summarized by Bonferroni confidence intervals (adjusting for all the 441 significant genes in at least one cluster). Dots represent the average of features values across samples.

Cluster 1’s signature was composed by genes mapped on chromosome 20. A group of 56 of the 57 genes exhibited significant copy loses in Cluster 1. Of this group, 50 genes (*ATRN, AP5S1, TMEM230, MGME1, NDUFAF5, CENPB, CRLS1, CRNKL1, CSNK2A1, DDRGK1, DSTN, DTD1, ESF1, FAM110A, FASTKD5, FKBP1A, IDH3B, ITPA, SMIM26, MAVS, MCM8, MKKS, MRPS26, NAA20, NOP56, NRSN2, NSFL1C, PANK2, PCNA, POLR3F, PSMF1, PTPRA, RBBP9, RBCK1, RRBP1, SIRPA, SMOX, SNPH, SNRPB2, SNRPB, SNX5, SOX12, STK35, TBC1D20, TRMT6, UBOX5, VPS16, ZCCHC3, ZNF133* and *ZNF343*) were also downregulated. From the group of genes with significant copy-losses and basal expression values (*TGM6, SOX13, PROKR2, PRND, OXT, LRRN4* and *FERMT1*), *LRRN4* and *FERMT1* were also significantly hyper- and hypo-methylated, respectively (Figure 3).

Cluster 4’s signature was composed by four genes mapping onto chromosome 6: *TDRD6*, *POLH*, *PAQR8* and *GUCA1A*. All these genes exhibited significant copy losses in Cluster 4, and all of them except *GUCA1A*, were also downregulated. Additionally, *POLH* was hypo-methylated, while *PAQR8* was hyper-methylated (Figure 3).

Cluster 6’s signature was composed by 23 genes mapping onto chromosome 11: *ALDH3B1, ANKRD13D, ANO1, AQP11, ARRB1, EMSY, CCND1, CTTN, KRTAP5-10, LRP5, LRRC32, TESMIN, MYO7A, NUMA1, PAK1, PPFIA1, RBM4, RPS6KB2, RSF1, SHANK2, TMEM134, TPCN2* and *USP35*. Every one of these genes exhibited significant copy gains, and all of them were also significantly upregulated, except for three genes with basal expression in Cluster 6: *MYO7A*, *LRRC32*, and *ALDH3B1*. Genes *USP35, SHANK2, MYO7A, LRRC32, CTTN, CCND1, ARRB1*, and *ALDH3B1* were additionally hypo-methylated, while genes *RSF1* and *PPFIA1* were hyper-methylated (Figure 4).

Cluster 7’s signature was composed by 24 genes mapping onto chromosome 6. All of these genes (*BTBD9, RRP36, CCND3, CNPY3, CUL7, FRS3, GUCA1A, BICRAL, KLC4, KLHDC3, LRFN2, MEA1, MED20, MRPL2, MRPS10, PEX6, PPP2R5D, RPL7L1, SRF, TAF8, TBCC, TOMM6, TRERF1*, and *UBR2*) exhibited significant copy gains. All of them were significantly up-regulated, except by *LRFN2, GUCA1A, BTBD9*, that had basal levels in Cluster 7. Genes *TRERF1, LRFN2*, and *FRS3* were additionally hypo-methylated, while *GUCA1A* was hyper-methylated (Figure 4).

Cluster 8’s signature was composed by 15 genes mapping onto chromosome 11. All of these genes (*ALDH3B1, ANO1, CCND1, CPT1A, CTTN, LRP5, MRPL21, NADSYN1, PPFIA1, RNF121, RSF1, SHANK2, TPCN2, UNC93B1*, and *USP35*) exhibited significant copy losses. All of them except *ANO1* (with basal levels in cluster 7) were significantly downregulated. Additionally, Genes USP35 and *NADSYN1* were significantly hyper-methylated, while *UNC93B1, RSF1, MRPL21* and *ANO1* were hypo-methylated (Figure 4).

## Discussion

Most pan-cancer classifications rely on molecular alterations that clearly discriminate between tissue of origin^13,15,16,19,20^. However, as soon as tissue effects were removed, we have found that the cancer signal immediately emerged. Distinct cancer classes were formed, containing tumors from different cancer types. These classes were also characterized by very specific functional groups of omic features. We also noticed that the main source of synergy across omics arose from strong positive correlations between gene expression and copy number events. The expression of regulatory elements within the group of selected features (including transcription factors and the micro RNA hsa-mir-615b) was, on the other hand, not associated with the expression of their predicted targets. These observations support the role of copy numbers as a major force affecting tumor progression^21–23^. Contrarily, methylation had a minor impact in the definition of clusters. This result could be due to the role of methylation in the determination and differentiation of cell types^24–26^. As a consequence, methylation effects were perhaps removed together with cell-type effects. Nevertheless, abnormal methylation patterns might still have had a role in the expression of some genes characterizing tumor classes (e.g. expression of *LRN4* and *GUCA1A* negatively correlated with promoter CpG islands average methylation).

The tumor clusters C1, C4, C6, C7, and C8 had exclusive signatures (i.e. different of every other cluster). Interestingly, the clusters without distinct individual signatures were the ones with more favorable outcomes (C3, C2, and C5). One possible explanation for this is the frequent correspondence between more dramatic molecular alterations and worse clinical outcomes^27,28^. To gain insights about possible biological interactions within each signature, we used the accompanying bibliographic results provided by the STRING database^29^ (see Material and Methods section). The literature suggests a wide overlap between signatures in terms of gene functions (cell growth, division, small RNA metabolism, protein synthesis, maturation and transport, and mitochondrial dysfunction). In the case of signature C1 (most genes down-regulated), the literature suggested *NOP56* (a core component of the small nucleolar ribonucleic protein) as a central element in the signature; interacting with *MKKS, NAA20* and *PTPRA* (genes with roles on mitotic division); *ESF1, SNRPB, SNRPB2, POLR3F* and *CRNKL1* (involved in small RNA processing), *PCNA* and *ITPA* (involved correct DNA replication and repair), *UBOX5, RRBP1*, *RBCK1* and *NRSN2* (protein synthesis, maturation and antigen presentation), *RBBPP9* (resistance to growth inhibition of TGF); *SIRPA* and *DSTN* (cell adhesion)^30–33^. In the signature C1, *NOP56* could be a candidate for future therapeutic intervention. Tumor suppressors *NRSN2* and *RBCK1* could also be considered.

The three downregulated genes from signature C4 were involved in small RNA maturation (*TDRD6*, micro RNA expression and maturation), cell proliferation (*PAQR8*, plasma membrane progesterone receptor), and DNA repair (*POLH*, DNA polymerase involved in DNA repair). From these groups, *PAQR8* and *TDRD6* could represent potential targets of therapy. Although neither of them has been directly related to cancer, other members of the PAQR family of progesterone receptors are known tumor suppressors, while *TDRD6* has been reported as frequently down-regulated in breast cancer, suggesting its potential use as biomarker^34^. In the case of signature C6 (most genes upregulated), the literature suggests *CTTN* as interacting with two groups of genes within the signature, either by co-expression or co-localization in amplicons. One group consisted of invasion and anti-apoptotic related genes (e.g. *SHANK*, *PAK1*, *PPFIA1*) and ion transport (*ANO1* and *TPCN2*)^35,36^. The other group consisted of *CCND1* (cell cycle check points), *LRPS* (protein synthesis), *RSF1* (chromatin remodeling), and *USP35* (protein turnover; through amplicon-mediated overexpression in breast and gynecological cancers)^37,38^. Patients with signature C6 could perhaps benefit by *ANO1* inhibitory therapy^36^.

Signature C7 was characterized by multiple genes co-expressing with *KLHDC3* (involved in homologous recombination): *MEA1* (spermatogenesis), *CNPY3* (protein folding, antigen presentation), *PPP2R5D* (direct catalytic activity), *RRP36* (small RNA synthesis), *CCND3* (cyclin, cell cycle checks points), and *MED20* (transcription). *KLHDC3* also belongs to the protein turnover and antigen presentation pathway, together with *CUL7* and *UBR2*. The literature also suggests another group of co-expressing genes within signature C7, consisting of *RPL7L* (ribosome), *MRPL2* and *MRPS10* (mitochondrial ribosome). These genes have also been found to physically interact in cell culture^39,40^. Signature C8 genes remarkably overlapped with signature C6 genes, but exhibited opposite regulation (i.e. up-instead of down-regulated). Additionally, the literature suggests interaction between *CCND1*, *NADSYN1* and *MRPL20* in signature C8^41,42^. *NADSYN1* has been proposed as target of inhibitory therapy in cancer^43^, while *MRPL20* has been suggested as biomarker for gastric cancers^44,45^. From symmetry with signature C6, patients with signature C8 might possibly benefit from *ANO1* inhibitory therapy^36^.

The molecular classification of tumors generated clusters with clear differences in prognosis and severity, with C3 exhibiting better outcomes than the remaining clusters. C3 also resembled a previously reported “inflammatory” type, in terms of immune infiltration and cancer type composition (enriched for prostate adenocarcinoma, thyroid, and pancreatic carcinomas and having elevated values of markers for CD4+ Th17 and Th1 cells and low genomic instability)^18^. Although the remaining clusters were clearly distinguished in terms of altered molecular processes, they were highly similar in terms of clinical and demographic characteristics. Further exploration of the link between clusters’ signatures and cancer phenotypes could aid in the development of novel biomarkers and therapies. For instance, signatures in clusters enriched for metastatic samples (C4 and C8) that remain one of the most severe cancer phenotypes could aid in the development of more efficient therapies^46,47^. Similarly, signatures could also rapidly address differences in tumor heterogeneity (e.g. C8 and C5 were notoriously more heterogeneous than the rest). Differences in immune infiltration (C6 with the lowest activated natural killers’ infiltration and C8 with the lowest lymphocytic one) could also imply the potential use of signatures to aid in immunotherapeutic decisions.

Given the possibility of unveiling different biological channels altered in tumors of similar clinical and molecular characteristics, we believe this novel pan-cancer classification could aid in the identification of therapies for cancers without standard of care.

## Supporting information

Supplementary Figures and Caption for Table S1

## Acknowledgments

All results shown here are in whole or part based upon data generated by the TCGA Research Network: https://www.cancer.gov/tcga.

## Material and Methods

### Pan-cancer data

The TCGA offers a demographically diverse sample with comprehensive and modern multi-omic data. We retrieved data from 5,408 from 33 cancer types made available by the Genome Data Commons (GDC) repository^48^, via the TCG-Assembler R package^49^. Omic data consisted of curated level-three data of genome-wide gene expression (GE), DNA methylation (METH), and copy number variants (CNV) profiles by tumor sample. GE profiles by sample corresponded with the logarithm of RNA-Seq counts by gene (Illumina HiSeq RNA V2 platform). METH profiles corresponded with CpG sites B-values from the Illumina HM450 platform, summarized at the CpG island level, using the maximum connectivity approach from the WGCNA R package^50^, and further transformed into M-values (M=β/(1-β); Du et al. 2010). CNV profiles corresponded to gene-level copy number intensity derived from Affymetrix SNP Array 6.0 platform, using human genome V19 as reference. The quality-control filtering process included the exclusion of features with all zeros, or coefficient of variation less than 1%. Samples or features with a disproportion of missing data (>20%) and/or single-sample batches were also excluded. Within the remaining samples, missing values were imputed by k-near neighbors, with k = 3. Each omic block was adjusted by batch effects using ComBat^52^. Final sample size after retaining subjects with information for all three omics was n=5,408.

Demographic information included gender, self-reported race and ethnicity, and patient’s age at the moment diagnosis (Table 1). Clinical information consisted of overall survival time and vital status at the final follow up, type of sample (from primary tumor, metastases, or normal tissue), tumor free fraction. We also used previously information from “The Immune Landscape of Cancer”^18^ with significant differences between clusters according to Kruskal-Wallis tests^53^. These covariates included: intra-tumor heterogeneity fraction (as subclonal genome fraction), and rates of non-silent mutations, aneuploidy, homologous recombination defects (all three derived as deviations from the normal genome), proliferation (normalized difference between number of dividing and non-dividing cells), and information from immune infiltrations (including scores for CD4+ cells, macrophages, lymphocytes, and natural killers) (See supplementary material in ^18^ for a detailed description of the scores calculation). Briefly, immune infiltration fractions were derived by CIBERSORT^54^, assigned to different cell classes, and multiplied by the leukocyte fraction derived from methylation data^18^.

### Omic integration, clustering and features selection

Our method can be conceptually described by the following four steps.

#### Step 1) Identification of major axes of variation and features selection

Integrative methods should be able to capture combined effects across omic sites that could either span across omic layers (e.g. epigenetics, gene expression, etc.) or extend genome wide (e.g. considering concomitantly contiguous CpG sites or even separated away sites). Let,

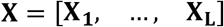

where **X**_*l*_ *l:{1,…, L}* is a matrix representing the **l**-^th^ omic, which row *i^th^* contains information representing a sample on one subject, and column *j^th^* represents an omic feature (e.g., a feature could be the expression of a specific gene, or the methylation level for a given CpG site). Each group of features coming from a different omic block is centered, standardized, and divided by 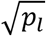 where *p_l_* is the number of features from the **l**-th omic block. This is done so larger groups of features do not dominate the data integration step. Next, we conduct a sparse Singular Value Decomposition (**sSVD**) of **X** to generate one factor that collapses the redundancies in the omics (by creating independent columns representing the independent signals across omic features) and one that collapses redundancies across samples, grouping subjects with similar signaling. This linear factorization can be represented as **X = ZW**, where **Z** represents (linearly) independent axes of variability across subjects (i.e. a lower rank approximation), while **W** represents loadings representing the contribution of each omic feature to this variability. This representation is common to many unsupervised omic integration methods, but is independent of distributional assumptions on each element. In this formulation, **Z** and **W** can obtained by minimizing:

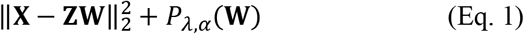

To the left of the plus sign is the Frobenius norm (a matrix analogous of Euclidean distance) of the difference between **X** and the product of **Z** and **W**. To the right of the plus sign is a penalty on the elements of **W** to impose sparsity. The purpose of this penalty is to zero-out those features with minor contributions to the columns of **Z**. To remove the effect of tissues, or other covariates that can influence the selection of features, we pre-multiplied **X** by **I** – **Q**(**Q’Q**)^−1^**Q’**, where **I** is a diagonal matrix of ones, and **Q** is an indicator matrix to represent the membership to a given organ or tissue.

#### Step 2) Identification of omic features (expression of genes, methylation intensities, copy gains/losses) influencing the axes

The linear decomposition achieved by SVD is an intuitive and straightforward way of integrating omics. However, the variability across omics can be governed by just a few features (i.e. highly *sparse* data) or by groups of interdependent features (i.e. very *redundant* data). To handle these limitations, we chose *P_λ, α_* **(W)** to be the Elastic Net penalty ^55^, 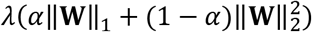, where *α* balances the regularization between LASSO and ridge regression types of regularization, and *λ* is associated with the degree of sparsity (i.e. how many features enter in the model?). Unlike LASSO, EN can select groups of correlated features, while zeroing out the irrelevant ones^56^. Equation 1 is solved by obtaining ***z_1_w_1_*** (where ***z_1_*** is the first column of **Z** and ***w_1_*** is the first row of **W**) with coordinate descent for given values of *λ* and *α*, following the algorithm of ^57^, as implemented in^58^, but with the following thresholding operator: sign(***w_1_***)| |***w_1_***| - *λα* |_+_ / *λ* (**1** – *α*) (where |***x***|_+_ represents the positive part ***x***). Consecutive layers are then obtained by subtracting the previous ones from **X** and repeating the same procedure, as many times as the number of desired axes of variation. The optimal value for *λ* was empirically determined, as suggested by^57^. We start by 1) calculating **W** over a dense grid of values for *λ* (lower *λ* yields less sparsity), 2) calculating the proportion of variance of **X** explained by **ZW** *(PVX)* for each *λ*, and 3) choosing the *λ* at which *PVX* has its minimum second derivative. Since *PVX* decreases monotonically with *λ*, this point represents a drastic drop on *PVX*, suggesting that the most relevant features accounting for the data variability are already incorporated^57^. The value *α* was fixed to 0.5 to have an equal contribution of LASSO and Ridge penalties. Once a subset of features was selected, we mapped them onto genes using annotation data of genomic position downloaded from the USCE web browser tool (GRCh38^59^). The enrichment of functional classes (ontologies, pathways, complexes, etc.) among these genes was tested using the Enrichr package^60^.

#### Step 3) Mapping major axes of variation via tSNE and cluster definition by DBSCAN

Additionally, SVD can be coupled with non-linear embedding methods to deal with highly heterogeneous data. Here, we applied *t* - Stochastic Neighbor Embedding (tSNE) on **Z**^14^. tSNE is a technique that efficiently takes on local neighborhoods present in high dimension (eventually representing clusters of data), and conserves them while projecting onto a lower dimensional display^61^. This makes tSNE a very powerful technique to reveal clusters, even in very heterogeneous and convoluted data settings^62^. The algorithm has two fundamental parameters: perplexity (which accounts for the effective number of local neighbors), and cost (related to the difference between the neighborhood’s distribution in the higher and lower dimensional spaces). Since low cost is an indication of displays more likely to reveal clusters, we selected the maps corresponding with the lowest costs among perplexities of 50 and 100, using 100 thousand iterations to ensure convergence. We applied Density-Based Spatial Clustering of Applications with Noise (DBSCAN^63^) to identify clusters. DBSCAN is one of the most powerful clustering techniques to delimit clusters of irregular shape, such as the ones tSNE produces^64^. Essentially, DBSCAN identifies groups of densely packed points, without the need of specifying the number of clusters a priori^63^. Neighborhoods of nearby points can then be tuned by evaluating different cluster partitions over a grid of possible neighborhood sizes. We tuned this parameter by maximizing the Silhouette score, as in Taskensen et al. 2016.

#### Step 4) Molecular and clinical characterization of clusters

The association between clusters and scores representing genes and functional classes selected, was studied to define the signatures representing each cluster. Scores were calculated by tacking the columns of **X** mapping onto a gene, or functional class, and post-multiplying it by the corresponding elements of **W’**. Due to the transformations of features values within each omic block (e.g. logarithm of standardized RPKM counts, Beta to M-values for CpG islands), scores can be considered to be approximately normal. Using the scores of each gene and functional class as response, and the clusters as explanatory variables, we then conducted a series of ANOVA tests to determine what genes or functional classes were significant in at least one cluster. All pairwise comparisons between significant genes and functional classes were studied via Tukey tests. Gene signatures were defined based on those genes significantly deregulated in a single cluster. For both types of tests, we used a Bonferroni multiple-test correction with P(type I –error) = 0.05 / {#selected genes and functional classes}.

To discuss the possibility of physical or functional relationships between the genes in each signature, we used the STRING data base of protein-protein interactions^29^. We considered an interaction as biologically meaningful whenever it was backed up by empirical data, such as immune precipitation, microarrays, curated databases, etc. Interactions suggested by text-mining (two genes reported in the same scientific publication) were not considered, except in the cases when a publication’s results gave evidence of interaction (e.g. genes co-expressing, co-locating, etc.).

The association between clusters and phenotypes (e.g. clinical, demographic, and immunologic covariates) was evaluated via Kruskal-Wallis test^53^ (non-parametric analogous of ANOVA). All significant results were further evaluated by Dunn test^65^ for pairwise differences (non-parametric analogous of Tukey tests). All steps of our method were implemented in the R programming language^66^, using irlba^67^, dbscan^63^, and Rtsne^68^ packages.

