## Supplementary Figures and Caption for Table S1 for "Multi-omic signatures identify pan-cancer classes of tumors beyond tissue of origin"

**Supplementary Information**

**Table S1: Complete list of genes significantly de-regulated in at least one pan-cancer cluster**. All the genes significantly different in at least one cluster are sorted by chromosome and genomic position. Significant differences between clusters are represented with different letters. Genes highlighted with red represent known cancer-related genes.

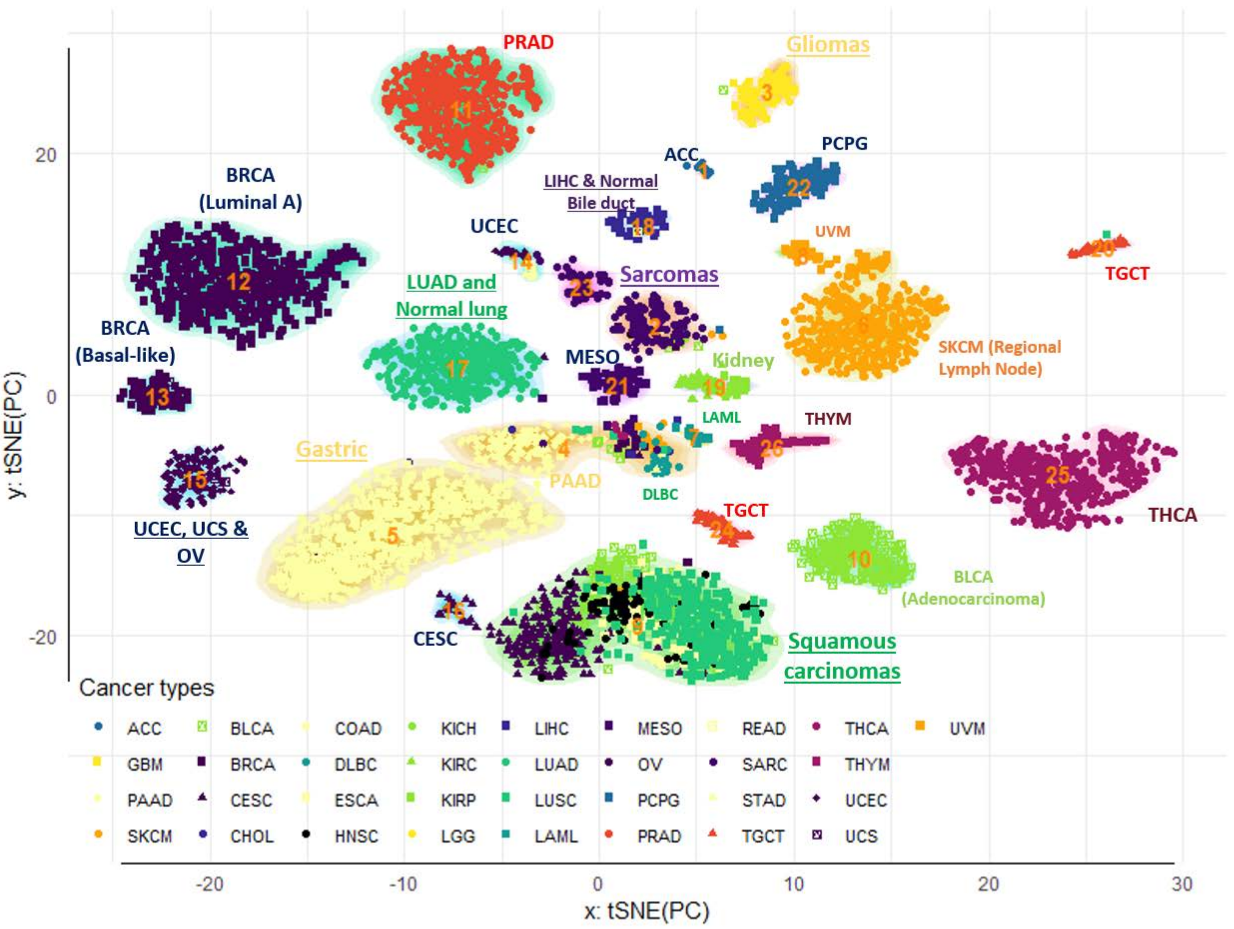


**Figure S1: Pan-cancer clustering of tumor samples (no tissue effects correction, no sparsity constrains on the features activities)**. Tumor clusters were obtained by sequential application of tSNE and DBSCAN algorithm for 5,408 samples from 33 cancer types. The contours reflect cluster membership, and the points' colors and shapes represent similar anatomical site and cancer type, respectively. The two dimensional tSNE projection was obtained from the first 50 principal axes of the extended omic matrix, after removing the first two. Extended omic matrix contained appended values of gene expression, DNA methylation, and copy number variant intensity. Integers represent individual clusters. Clusters were also annotated in terms of their most enriched histological/molecular subtypes.

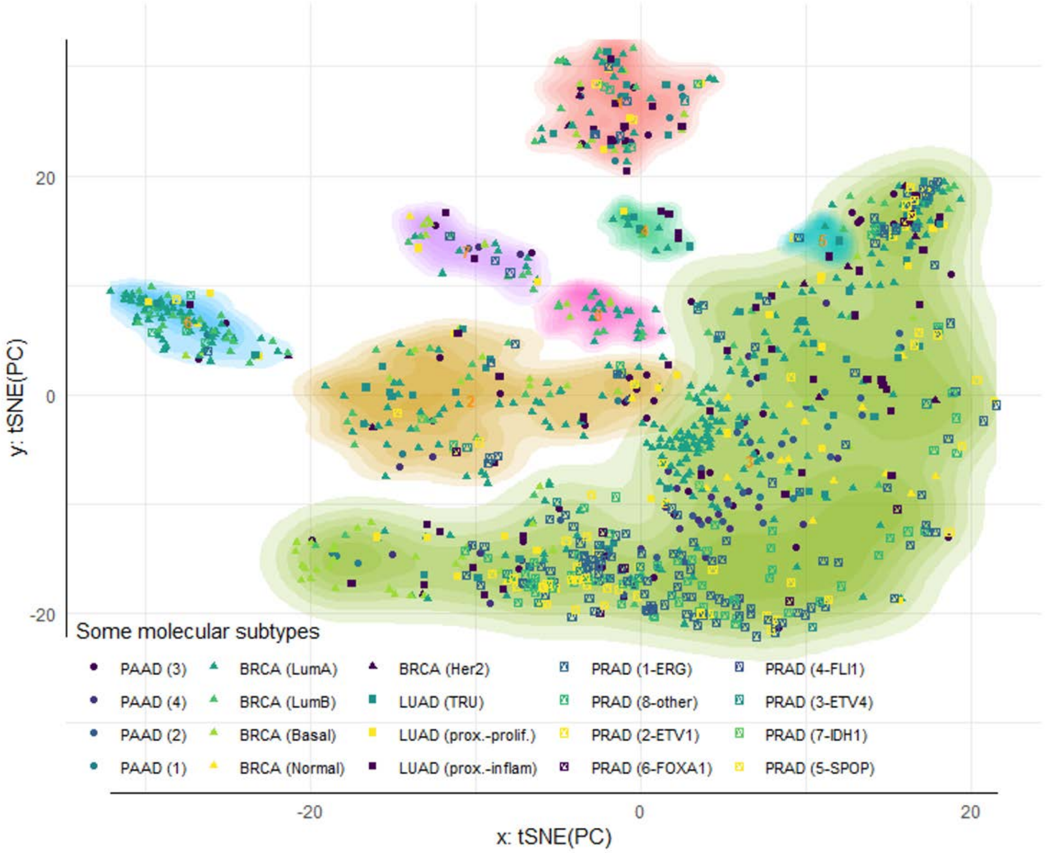


**Figure S2: Re-classification of tumors after removing tissue effects does not agree with previously reported molecular subtypes**.

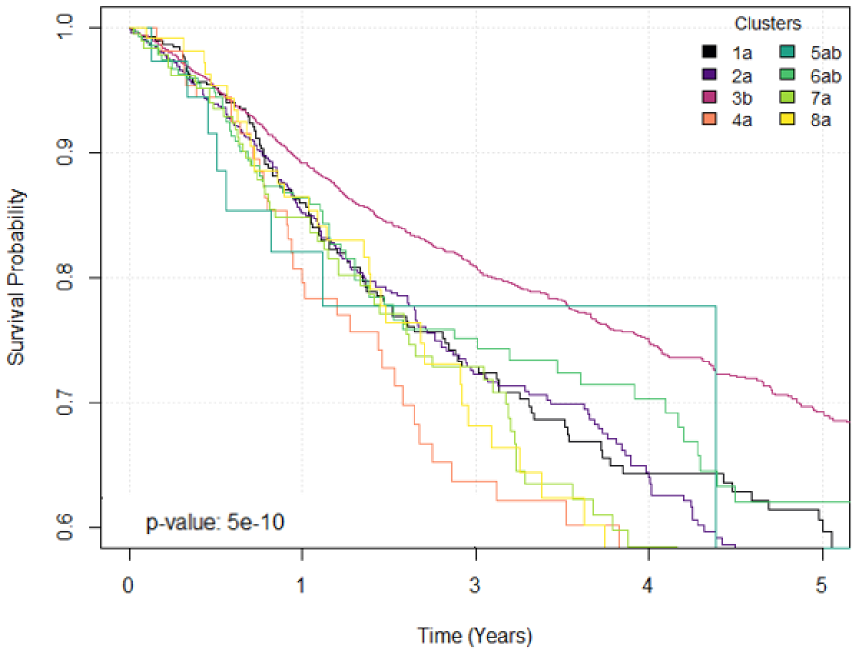


**Figure S3: Survival curves by pan-cancer tumor clusters**. The figure show Kaplan-Meier curves highlighting the survival probability by time in years for each cluster. Log-rank tests were performed to determined significant differences between curves. The legend shows the results of multiple comparison between survival curves. Statistical differences are represented with different letters.

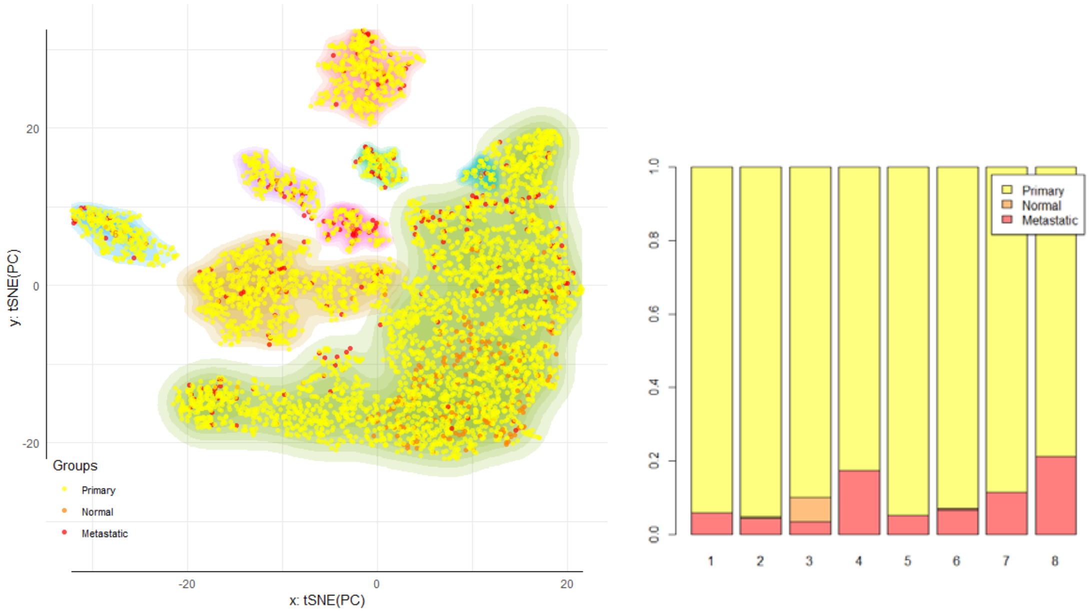

**Figure S4: Re-classification of pan-cancer tumors after removing tissue effects reveals differences in sample type composition**. The relative position and number of primary, normal, and metastatic tissue samples is shown. The figure at the left shows the samples location by clusters. The figure at the right shows the relative proportion of sample types by cluster.

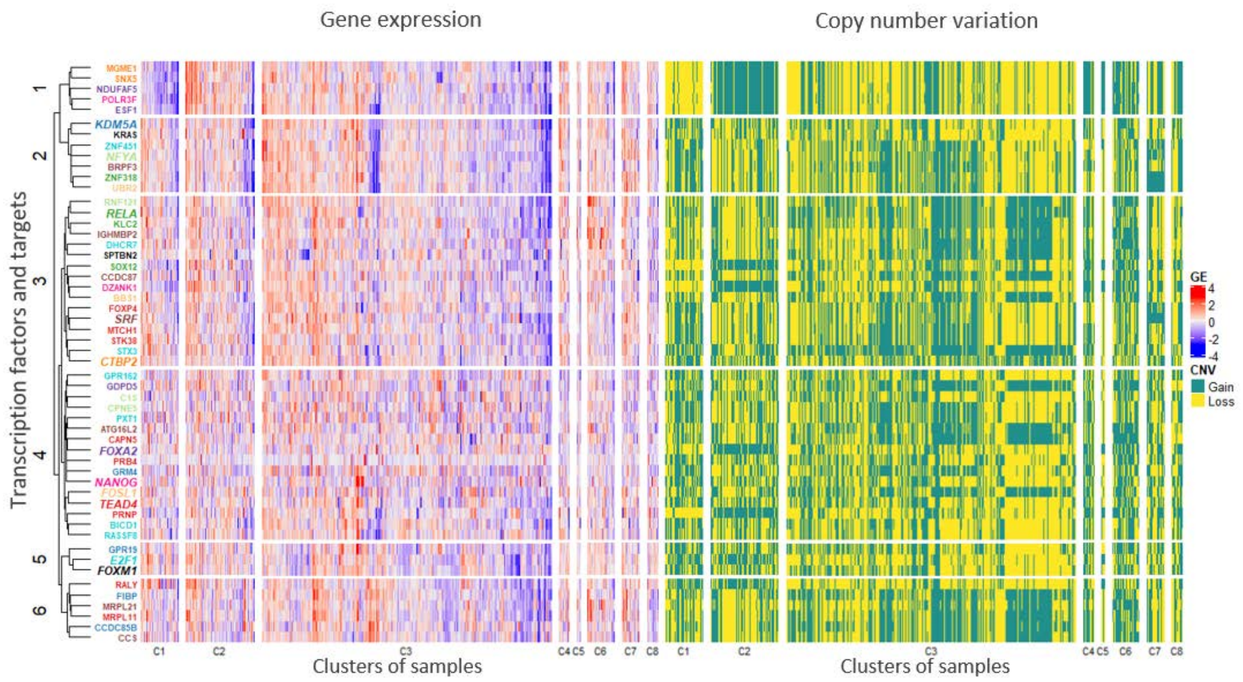


**Figure S5: Relationship between gene expression and copy number variation for transcription factors and their targets within the set of selected features**. The expression and copy number variation by gene is shown by cluster (C1-8). The colors by gene name represent groups defined by different transcription factors and their targets (e.g. black represents the group of *FOXM1* and its targets *KRAS* and *SPTBN2*). TFs names are shown with italic and larger font size. The number at the left of the dendogram represent grouping of genes based on k-means clustering.
